# A Model Visualization-based Approach for Insight into Waveforms and Spectra Learned by CNNs

**DOI:** 10.1101/2021.12.16.473028

**Authors:** Charles A. Ellis, Robyn L. Miller, Vince D. Calhoun

**Affiliations:** Tri-institutional Center for Translational Research in Neuroimaging and Data Science: Georgia State University, Georgia Institute of Technology, Emory University, Atlanta, GA 30303 USA; Tri-institutional Center for Translational Research in Neuroimaging and Data Science: Georgia State University, Georgia Institute of Technology, Emory University, Atlanta, GA 30303 USA.

## Abstract

Recent years have shown a growth in the application of deep learning architectures such as convolutional neural networks (CNNs), to electrophysiology analysis. However, using neural networks with raw time-series data makes explainability a significant challenge. Multiple explainability approaches have been developed for insight into the spectral features learned by CNNs from EEG. However, across electrophysiology modalities, and even within EEG, there are many unique waveforms of clinical relevance. Existing methods that provide insight into waveforms learned by CNNs are of questionable utility. In this study, we present a novel model visualization-based approach that analyzes the filters in the first convolutional layer of the network. To our knowledge, this is the first method focused on extracting explainable information from EEG waveforms learned by CNNs while also providing insight into the learned spectral features. We demonstrate the viability of our approach within the context of automated sleep stage classification, a well-characterized domain that can help validate our approach. We identify 3 subgroups of filters with distinct spectral properties, determine the relative importance of each group of filters, and identify several unique waveforms learned by the classifier that were vital to the classifier performance. Our approach represents a significant step forward in explainability for electrophysiology classifiers, which we also hope will be useful for providing insights in future studies.

**Clinical Relevance:** Our approach can assist with the development and validation of clinical time-series classifiers.

## I. Introduction

In recent years, deep learning models, like convolutional neural networks (CNN), have been increasingly applied to electrophysiology time-series data [1]–x[4]. Relative to previous approaches that used extracted features, these approaches are much less explainable [5], and novel explainability methods have been developed as a result [1], [6], [7]. Most of these methods provide insight into spectral features learned by classifiers, and while spectra are important features, specific waveforms are more useful in many electrophysiology modalities. In this study, we present the first method focused on explainability of EEG waveforms and spectra learned by convolutional neural networks (CNNs).

Prior to the growth of modern deep learning, electrophysiology studies involving machine learning often used extracted features. These extracted features sometimes summarized time domain properties and often included spectral activity, particularly in electrophysiology domains like electroencephalography (EEG) or magnetoencephalography (MEG) [5]. The use of extracted features enables existing interpretability and explainability approaches to be applied. However, as the role of deep learning in electrophysiology studies has further developed, classifiers have been increasingly applied to raw time-series data. While this enables the automated extraction of features, traditional explainability approaches often do not provide clear global results for classifiers trained on raw time-series.

As such, a growing number of studies have begun to develop approaches to provide insight into deep learning classifiers trained on raw electrophysiology time-series. Most of these studies provide insight into key spectral features learned by classifiers [1], [6]–[10], and some studies have sought to gain insight into the relative importance of different modalities in multimodal classification problems [11]. Lastly, only a couple studies have presented methods capable of providing insight into waveforms learned by classifiers [1], [8]. This is important because waveforms, not just spectra, are vital to classification in electrophysiology [12]. These studies do not provide adequate global approximations of the key waveforms learned by the classifier. Methods like those used in [1] would require the manual inspection of importance results for each sample in the dataset to gain an understanding of key waveforms, and while [8] does provide insight into waveforms, it primarily provides insight into the combinations of spectra that maximize the activation of a model.

Visualizing model weights provides an alternative to existing attempts at insight into key waveforms learned by CNNs from electrophysiology data. Many existing CNN architectures use short filters in their first layer [13], which makes it difficult to visualize the filters and understand the extracted features. In contrast, one EEG study used longer filters that can be visualized for insight into key features [9]. However, [9] only analyzed the spectral features learned by the model.

In this study, we propose a novel model visualization-based approach that provides the first global insights into the waveforms learned by classifiers. Our approach also provides insight into the spectral features learned by the classifier and into the importance of each of the filters in the first layer of the classifier. We demonstrate the viability of our approach within the context of sleep stage classification, a well-characterized domain, and find that our results conform to domain knowledge.

## II. Methods

Here we describe our study approach.

### A. Description of Dataset

We use the Sleep Cassette data from the Sleep-EDF Expanded dataset [14] on PhysioNet [15]. The dataset has 153 20-hour recordings from 78 study participants. The data was recorded at a sampling frequency of 100 Hertz (Hz), and we used the FPz-Cz electrode. The data was assigned to Awake, Rapid Eye Movement (REM), Non-REM 1 (NREM1), NREM2, NREM3, and NREM4 stages in 30-second intervals.

### B. Description of Data Preprocessing

We separated the data into 30-second segments using their annotated classes. We removed the Awake data at the start of the recordings and many samples at the end of the recordings to alleviate class imbalances. We also combined NREM3 and NREM4 into a single NREM3 class based on clinical guidelines [12] and z-scored each recording. Our final dataset had 85,034, 21,522, 69,132, 13,039, and 25,835 Awake, NREM1, NREM2, NREM3, and REM samples, respectively.

### B. Classifier Development

We adapted the classifier initially developed in [9]. The architecture was originally developed for segments longer than 30 seconds, so it was necessary for us to update the architecture for our data. Our architecture is shown in Figure 1. We used 10-fold cross-validation when developing the classifier, randomly assigning all samples from 63, 7, and 8 participants to the training, validation, and test groups, respectively. We accounted for class imbalances via a class-weighted categorical cross entropy loss function, and we used stochastic gradient descent with an initial learning rate of 0.015 that decreased by 10% if 5 training epochs passed without an increase in validation accuracy. Additionally, we used early stopping if the classifier went 20 epochs without an improvement in validation accuracy with a maximum of 30 epochs. We also used a batch size of 100. To assess the classifier performance, we computed the recall, precision, and F1 score for the test data across each fold and subsequently computed the mean and standard deviation of each metric.

**Figure 1.**
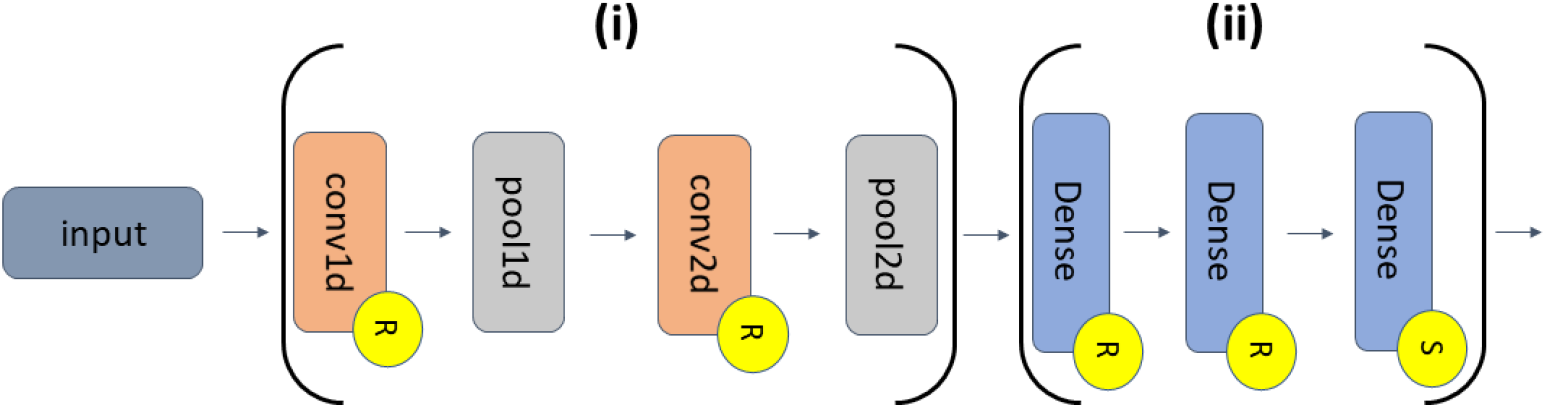
CNN Architecture. Sections (i) and (ii) indicate the feature extraction and classifier portions of the CNN, respectively. (i) consists of a 1D convolutional (conv1d) layer (30 filters of length 200 and stride of 1), a 1D max pooling (pool1d) layer (pooling size of 15 and stride of 10), a conv2d layer (400 filters of 30 x 25 dimensions and stride of 1 x 1), and a pool2d layer (pooling size of 10 x 1 and a stride of 1 x 2). (ii) consists of 2 dense layers (500 nodes and a final dense layer with 5 nodes. Layers with an “R” or an “S” indicate that they have ReLU or softmax activation functions, respectively.

### C. Explainability – Spectral Clustering and Filter Importance

A previous study visualized the spectral features learned by the classifier [9]. To simplify the visualization and better identify overall patterns learned by the classifier, we performed a fast Fourier transform on each of the filters, calculated the power in 1 Hz intervals between 0 Hz and 50 Hz, and performed k-means clustering on the power values. It should be noted that across all of our explainability analyses, we used the test data and model weights from the fold with the highest weighted F1 score. We performed an initial round of clustering with the silhouette method to determine the optimal number of clusters between 2 and 15. After finding the optimal number of clusters, we reclustered with the optimal number. We used 50 initializations in our initial clustering and 100 initializations when using the optimal number of clusters.

After clustering, we applied layer-wise relevance propagation (LRP) to determine the importance of each filter. LRP has been applied to electrophysiology analyses previously [16]. LRP does not ordinarily indicate the relative importance of model filters. However, it provides a more methodologically sound alternative to the perturbation approach in [9]. In order to apply the method, we calculated the LRP relevance for the first layer activations of all test samples. We specifically used the αβ-rule with an α of 1 and a β of 0, which allowed us to only examine positive relevance. After determining the total relevance of the activations associated with each filter, we computed the average within-cluster filter activation for samples in each class.

### D. Explainability – Filter Perturbation Analysis

While our first analysis gave insight into the spectral features in each filter and the relative importance of those filters, the key innovation of our study is our filter perturbation analysis. In this analysis, we iteratively perturbed the weights of the conv1d layer using a sliding window with filter padding that assigned values of zero to weights within the window. After each perturbation, we calculated the percent change in the weighted F1 score of the classifier following perturbation. We used a sliding window with a length of 25 and stride of 1. While we examined other window lengths, we excluded them in this manuscript due to space constraints.

## III. Results AND Discussion

In this section, we describe and discuss our analysis results.

### A. Classification Performance

Table 1 shows our model classification performance results. The classifier had the highest performance for the Awake class across all metrics, possibly due to the presence of high amplitude artifact present in some Awake samples [8]. After Awake, NREM2 had the second highest performance across all metrics. NREM3 and REM performance were relatively high. In contrast, NREM1 performance was low but consistent with previous studies that have also found classifying NREM1 samples difficult [17].

**TABLE I.**
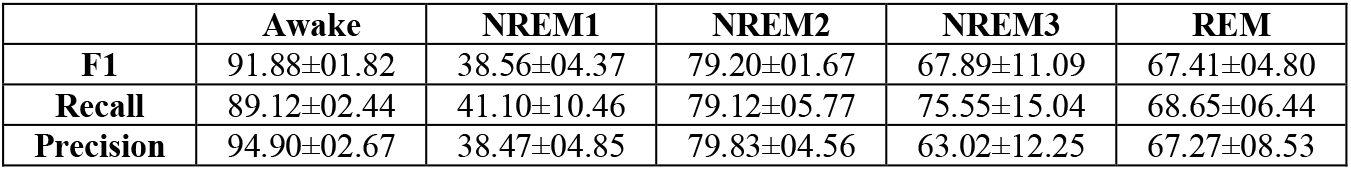
Classification Performance Results.

### B. Spectral Clustering and Filter Importance

Our spectral clustering analysis found that using 3 clusters was optimal. Figure 2 shows the spectral values of the filters in each cluster. Cluster 1 was the largest cluster, with 16 filters. It extracted large amounts of upper θ (4 – 8 Hz), lower α (8 – 12 Hz), mid to high-range β (12 – 25 Hz), and γ (25 – 50 Hz) activity. Cluster 2 was the second largest cluster with 11 filters, and it extracted δ (0 – 4 Hz) and θ activity. Lastly, cluster 0, which had 3 filters, extracted α and lower β-band activity.

**Figure 2.**
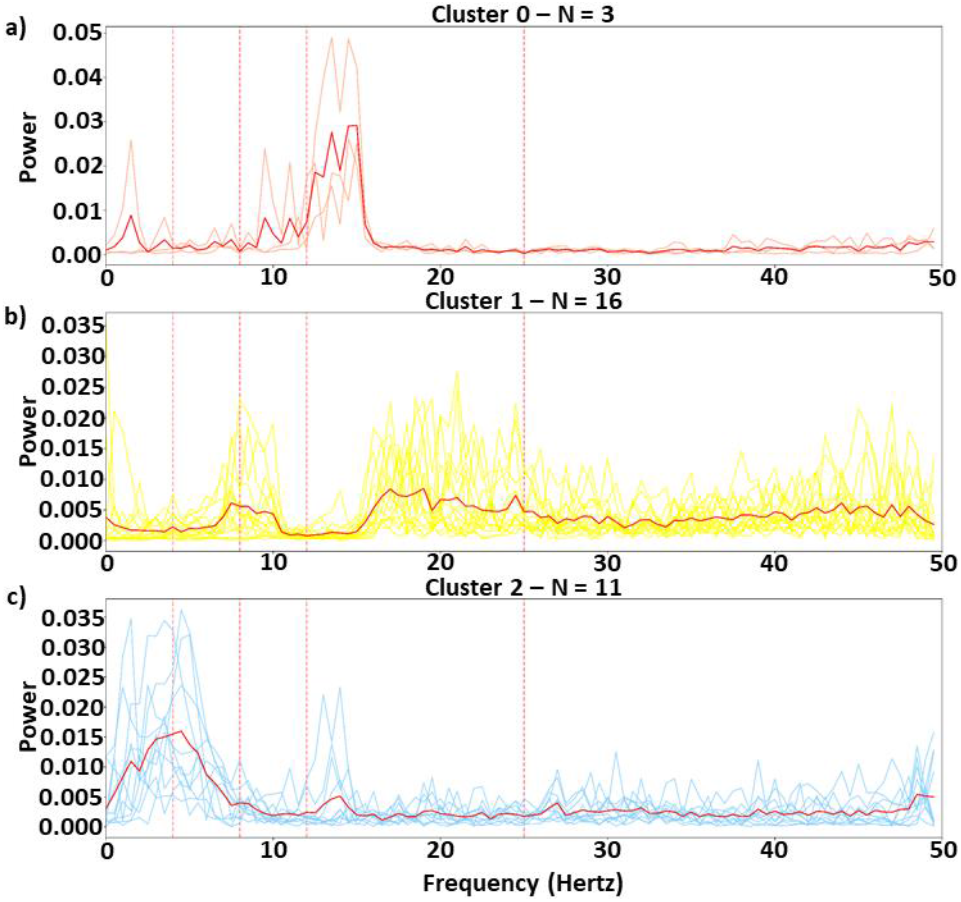
Filter Spectra. Panels a, b, and c show the spectra of all filters in clusters 0, 1, and 2, respectively. The x-axis shows the frequency in Hz, and the y-axis indicates power. The thick red line in each panel is the cluster center. The vertical dashed lines indicate the boundaries of the canonical frequency bands (i.e., δ, θ, α, β, and γ). Note that values for clusters 0, 1, and 2 are light red, yellow, and light blue, respectively.

Figure 3 shows our LRP filter importance results. Overall, cluster 2 demonstrated much more relevance than the other clusters and a particular emphasis on NREM3. NREM3 is characterized by δ [12], which was one of the main frequencies extracted by cluster 2. Cluster 0 was most relevant to NREM2, which makes sense given that sleep spindles are important to NREM2 and are found within the frequencies extracted by the cluster [12]. Interestingly, cluster 1 extracted θ and was most important to NREM1, which is characterized by θ [12].

**Figure 3.**
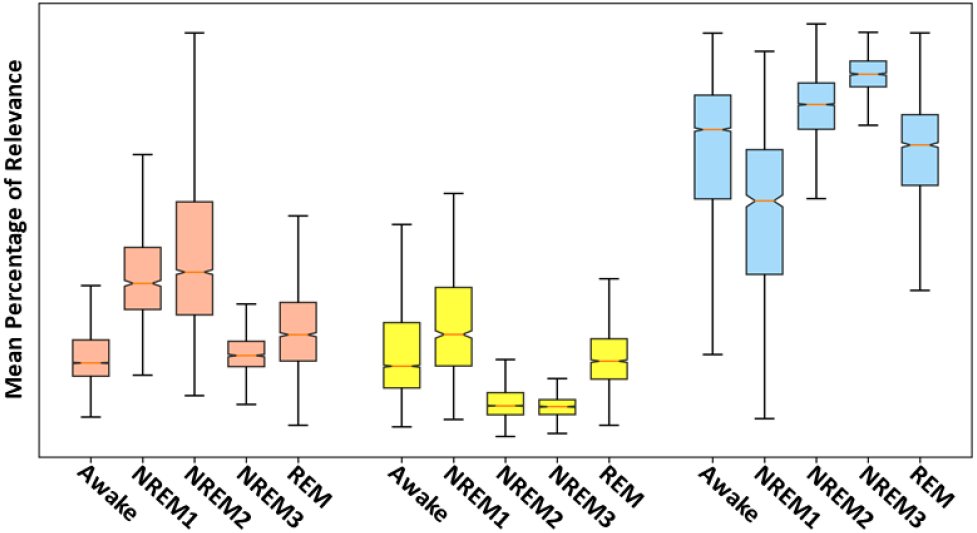
LRP Results. This shows the mean percent of filter relevance for correctly classified samples within each class and cluster. Clusters 0, 1, and 2 are light red, yellow, and light blue, respectively.

### C. Filter Perturbation Analysis

Figure 4 shows our results for the filter perturbation analysis. Interestingly, while the perturbation of most of the filter weights had relatively little impact upon the classifier performance, the perturbation of some sets of filter weights had large effects upon the classifier performance. Cluster 2 weights seemed to most significantly affect the classifier. The perturbation of the unique waveforms in filters 21 and 26 caused the largest change in performance across all filters. The waveform in filter 26 between 1 and 1.25 seconds is possibly an inverted k-complex, and the key waveform in filter 21 is possibly a vertex sharp wave. K-complexes and vertex sharp waves are associated with NREM1 and NREM2, respectively [12]. Additionally, perturbation of lower frequency waveforms (e.g., filters 20 and 29) impacted classifier performance. Filter 29, in particular, had δ waves between 0 and 0.75 seconds that are associated with NREM3 [12]. In contrast, perturbation of filter weights in clusters 0 and 1, which were characterized by higher frequency activity, seemed to have less impact upon the classifier. This likely occurred because the classifier learned more high frequency activity and was able to adapt to the perturbation of short windows. For example, filters in cluster 1 may have had higher frequency sleep spindles. In contrast, fewer cluster 0 and 1 weights had the low frequency activity and unique waveforms found in cluster 2, so the classifier could not adapt to the perturbation of those features.

**Figure 4.**
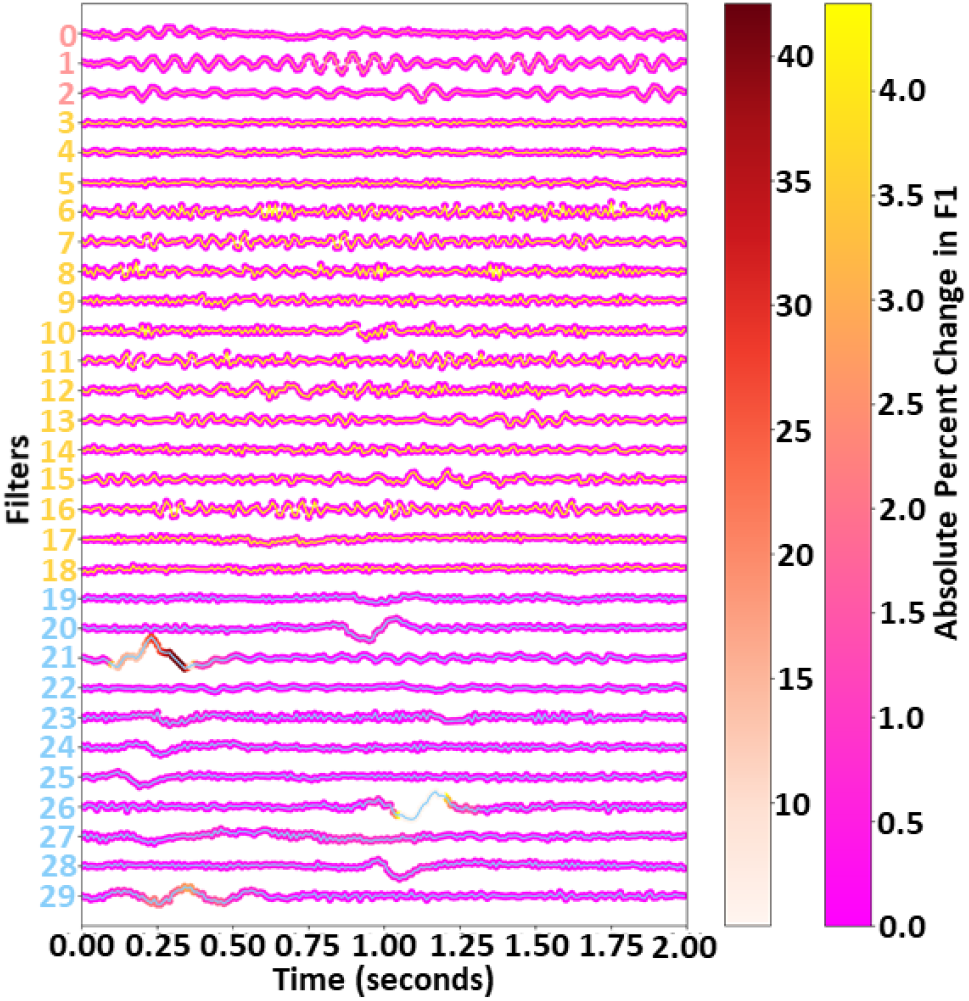
Filters and Perturbation Results. The x-axis shows the time point associated with each filter, and the filters are arranged on the y-axis. Cluster 0, 1, and 2 filter numbers are in light red, yellow, and light blue, respectively. To better visualize the heavy concentration of values between 0 to 5%, the two heatmaps indicate the absolute percent change in the weighted F1 score at each point. The leftmost and rightmost color bars cover values between 5 – 40%, and between 0 – 5%, respectively.

### D. Limitations and Future Work

In our filter perturbation analysis, we examined the effect of perturbation upon the weighted F1 score. Using class-specific F1 scores could indicate whether some features have class-specific importance. Additionally, we only included results for a window length of 25 in this study. In the future, we would like to more exhaustively investigate the effects of the window length parameter. Future studies may consider perturbing the filter spectra to better understand the relative importance of each spectral feature in the filters to the final classifier performance. Lastly, our clustering with filter spectra, instead of waveforms, greatly simplified the clustering but assumed that the key information captured by the filters was related to spectral power. Our choice was due to the inherent difficulty of clustering time-series based on features that can occur anywhere in the time course. Methods like convolutional autoencoders might help capture key features and be paired with traditional clustering approaches to for waveform-based clustering. Nevertheless, spectral clustering was not the main contribution of our work, and it did produce clearly defined groups of filters.

## IV. Conclusion

In this study, we present a novel model visualization-based explainability approach that provides key insights into waveforms learned by CNNs that preexisting approaches are incapable of providing. We train a CNN for automated sleep stage classification, a well-characterized domain, to demonstrate the reliability of the approach. We identify clusters of filters in the conv1d layer of the CNN that extract highly distinct spectral activity, and we identify the importance of each filter cluster to the identification of each sleep stage. Most importantly, we then apply a novel filter perturbation approach to identify specific waveforms in each filter that impact the classification performance, possibly identifying k-complexes and vertex spikes. This high degree of insight would be impossible without our use of long filters in the first layer, which distinguishes it from most existing electrophysiology classifiers. Our novel approach represents a significant step forward in explainability for electrophysiology classifiers, and we hope it will provide many novel insights in future studies.

